# Biohybrid tendons enhance the power-to-weight ratio and modularity of muscle-powered robots

**DOI:** 10.1101/2025.07.10.664167

**Authors:** Nicolas Castro, Maheera Bawa, Bastien Aymon, Sarah Wu, Annika Marschner, Sonika Kohli, Laura Rosado, Martin Culpepper, Xuanhe Zhao, Ritu Raman

## Abstract

Biohybrid robots powered by tissue engineered skeletal muscle have historically relied on architectures in which muscle actuators are placed directly on skeletons, thus limiting the accessible design space for such machines. By contrast, native musculoskeletal architecture relies on tendons to bridge the interface between muscles and skeletons, enabling precise, space-efficient, and energy-efficient force transmission. In this study, we use a mathematical model of the muscle-tendon-skeleton interface to design a biohybrid muscle-tendon unit composed of tissue engineered muscle coupled to adhesive tough hydrogel tendons. We show how tuning tendon stiffness and pre-tension modulates actuator performance, measure fatigue characteristics of our actuators over >7000 cycles, and tune skeleton stiffness to increase force transmission muscles to skeletons by ∼29X. Furthermore, we demonstrate an ∼11X improvement in power-to-weight ratio of muscle-tendon units as compared to previous demonstrations of robots powered by muscles alone. This work validates a robust approach for designing, manufacturing, and deploying muscle-tendon actuators that promises to enhance the modularity and efficiency of biohybrid robots.

## 1. Introduction

Biohybrid robots powered by tissue engineered skeletal and cardiac muscle have recently emerged as a robust approach for endowing machines with the adaptive capabilities of living tissues.^[1–3]^ Demonstrations of muscle-powered walkers,^[4–10]^ swimmers,^[11–17]^ pumps,^[18–21]^ grippers^[22–25]^, and mechanical computers^[26–28]^ have showcased the versatility of these actuators for powering a range of useful functions. Recently, integration of bioelectronic control and neural control of muscle actuators, coupled with efforts to incorporate real-time sensory feedback, has advanced the complexity and programmability of biohybrid machines even further.^[29–35]^ Moreover, studies of adaptation in muscle-powered robots have highlighted their ability to tune their morphology and function in response to changing environmental stimuli, such as exercise or injury,^[36–40]^ and generate multi degree-of-freedom motion.^[41]^ Overall, these multidisciplinary advances highlight the rapid growth of this field over the past decade, while also highlighting open challenges that must be addressed before living muscle actuators can be deployed to address real-world problems.

Biohybrid robots that require on/off control, exercise-mediated performance adaptation, and resilience to injury typically rely on skeletal muscle rather than cardiac muscle, as skeletal muscle is intrinsically capable of these behaviors and thus powers all voluntary movement in our bodies and many other living creatures (**Figure 1**A).^[42,43]^ Demonstrations of skeletal-muscle powered robots have primarily relied on placing a strip or ring of engineered muscle tissue directly on a flexible skeleton, such that linear actuation of the muscle drives skeleton motion along a defined direction.^[4–8,22,24,25]^ The robustness of the muscle-skeleton physical connection is dependent on interfacial friction between the muscle and skeleton and the fracture strength of the muscle. As a result, typical biohybrid robot designs require that a large percentage of the muscle is used to perform a wasteful structural role rather than an actuation role (Figure 1B).

**Figure 1.**
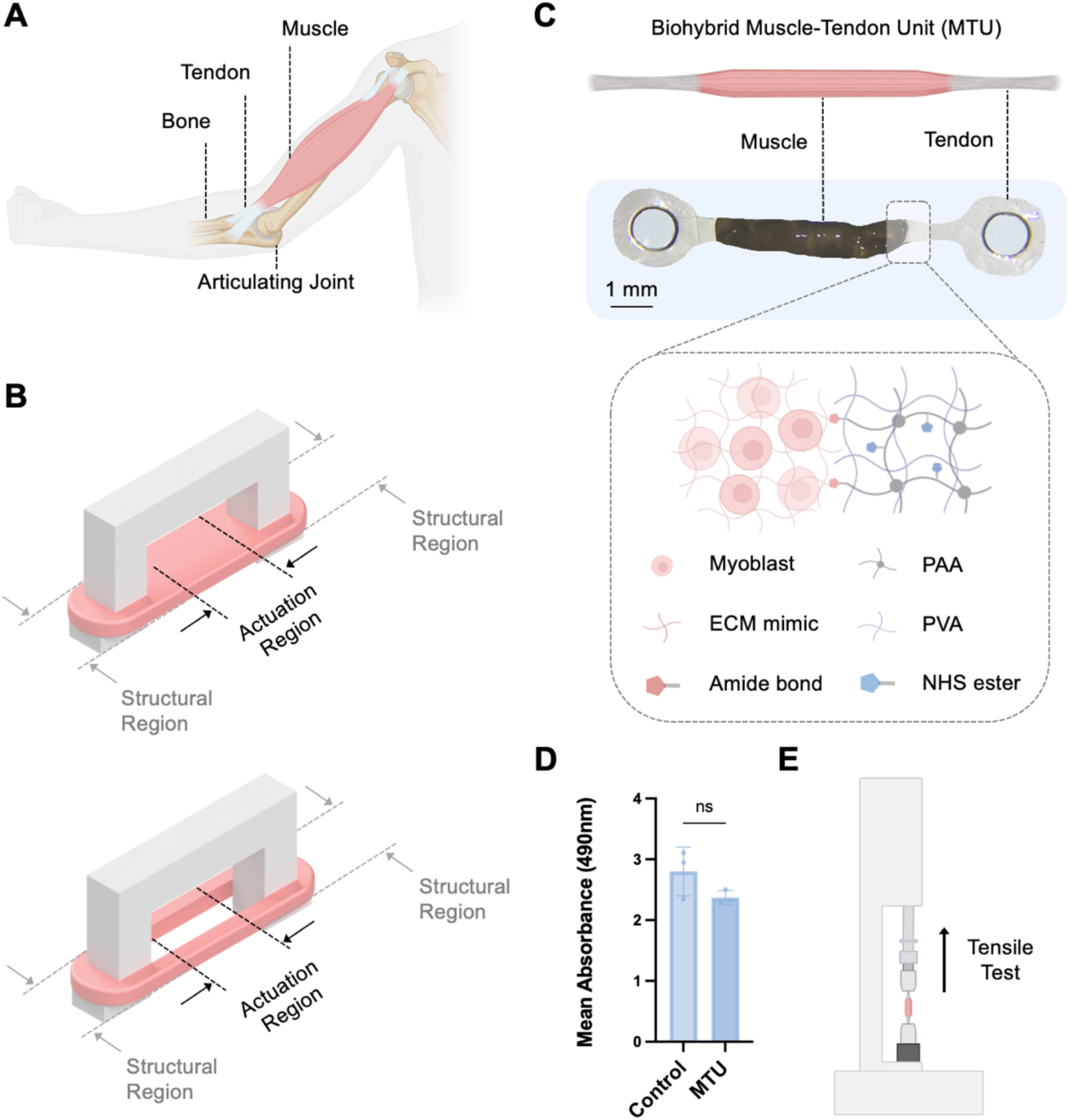
Biohybrid MTU. (**A**) Schematic of native muscle-tendon-bone architecture. Muscle contraction generates movement across an articulating joint. (**B**) Schematic of typical muscle strip (top) and ring (bottom) designs typically used for biohybrid robots. A large percentage of the tissue is typically used to form a robust connection with the skeleton and thus not used for actuation. (**C**) Biohybrid MTUs are composed of hydrogel tendon adhered to either end of a strip of tissue engineered skeletal muscle. The tendon is composed of an NHS-functionalized double network PVA:PAA hydrogel that forms covalent amide bonds with the engineered tissue formulated from myoblasts embedded in an extracellular matrix (ECM) mimicking hydrogel composed of 4 mg mL^−1^ fibrin and 30% v/v Matrigel. (**D**) Colorimetric quantitative viability assay demonstrating no significant difference in mean absorbance, representing viability, of muscle cultured alone (control) or in co-culture with PVA:PAA tendon (MTU). Welch’s t test, n = 3 per group. (**E**) Schematic of mechanical testing of biohybrid MTUs. Tests demonstrated that 1*x*1 mm muscle-tendon interfaces could withstand forces > 5 mN without delamination.

The portion of the muscle that is not being used for structural support generates useful contractile forces in response to electrical stimulation. Controlling the magnitude of the muscle’s contractile stroke, as well as the force transmitted from the muscle to the skeleton, requires tuning the skeleton’s stiffness: i.e. compliant skeletons yield large deformation and low transmitted force, while stiff skeletons yield low deformation and large transmitted force. To date, skeletal muscle-actuated machines have primarily focused on demonstrations of highly compliant skeletons that enable visualizing large deformations in response to stimulation, while limiting force transmission.

The first generations of biohybrid robots achieved compliance by fabricating skeletons entirely from soft polymers such as poly (ethylene glycol) diacrylate or polydimethylsiloxane (elastic moduli in the kPa to single MPa range, mimicking soft tissues)^[5,7,18]^ However, the effective stiffness of a skeleton fabricated from such compliant materials is often highly sensitive to changes in muscle placement and changes in stimulation frequency, thus limiting inter-device reproducibility.^[44]^ To address this drawback, a few studies have shown that skeletons manufactured entirely from rigid materials (elastic moduli in the GPa range, mimicking bone) can be made compliant via integrating flexure-based or articulating joints, thus improving inter-device reproducibility even at high-frequency operation while still enabling large deformations.^[22,24,44]^ While promising, such approaches still require coupling muscles to highly compliant mechanisms, in order to prevent nucleation of stress concentrations at the soft muscle-rigid skeleton interface that could lead to tissue damage and tearing.

Preventing stress concentrations at the muscle-skeleton interface thus remains a major challenge in biohybrid robotics, and has largely been addressed by: 1) increasing the percentage of tissue used for structural interfacial support and 2) constraining the upper limit of skeleton stiffness. By thus limiting biohybrid robots to wasteful designs and compliant mechanisms that yield low force transmission, the current state-of-the-art reduces the accessible design space and useful real-world applications of such machines.

Addressing this fundamental challenge in biohybrid robot design requires rethinking the paradigm of placing muscles directly on skeletons. This problem has already been solved in nature by the muscle-tendon unit (MTU) observed in vertebrate animals, in which muscles are physically connected to skeletons via elastic non-contractile tendons, enabling efficient force transmission from the contractile muscle to the bone.^[45,46]^ Tendons are composed primarily of linearly aligned collagen fibrils, yielding a uniaxially stiff tissue (elastic modulus range of ∼100s of MPa) that enables efficiently transmitting force from muscle (∼100s of kPa) to bones (∼10s GPa).^[47–50]^ This biological design has many key advantages including: 1) preventing muscle injury by providing a stiffness gradient from soft muscles to rigid bones that limits formation of stress concentrations at the interface of mechanically mismatched tissues; 2) providing space-efficient connections from bulky muscles to bones via a thin and flexible cable-like tissue that can wrap around articulating joints without being damaged; 3) enabling precise motions along many degrees of freedom by controlling the exact location and angle at which muscles attach to skeletons

In this study, we take inspiration from the native MTU to design a biohybrid MTU composed of a linear skeletal muscle actuator with tough hydrogel tendons at each end. The modular design of our biohybrid MTU enables coupling it to different skeleton designs to efficiently transmit force and power useful functions. We first demonstrate that tough adhesive hydrogel “tendons” can form robust bonds with tissue engineered muscle without compromising tissue viability or contractility. We then construct a mathematical model of muscle-tendon-skeleton interaction and investigate how tendon stiffness and pre-tension can be optimized to enhance actuator contractility. We also conduct fatigue tests across a range of stimulation frequencies to show that the muscle-tendon interface in MTUs is highly durable over > 7000 contraction cycles. While previous demonstrations of biohybrid robots have coupled muscle actuators to highly compliant skeletons in order to amplify motion, such designs inherently limit the force transmission capacity of these tissues since minimal effort is required to actuate a low-stiffness mechanism. By contrast, we leverage the modularity of our biohybrid MTUs to power a high-stiffness gripper skeleton, and show that increasing mechanism stiffness by 50X enables increasing the transmitted specific force of muscle actuators by ∼29X. The efficient design of biohybrid MTUs enables enhancing the power-to-weight ratio of our actuators by ∼11X as compared to previous biohybrid robots using the same muscle formulation. Taken together, these advances highlight the potential for a new generation of biohybrid MTU actuators that take inspiration from the native musculoskeletal interface to enhance the efficiency and modularity of biohybrid robots.

## 2. Results

### 2.1. Designing and manufacturing a modular biohybrid muscle tendon unit (MTU)

Prior efforts to replicate MTU architecture *in vitro* have focused on tissue engineering the myotendinous junction by culturing muscle and tendon cells in different regions of a hydrogel scaffold with a stiffness gradient.^[51]^ Some reports have also investigated the use of acellular polymeric tendons as anchors for tissue engineered skeletal muscle, enabling dynamic mechanical stretching of the muscle.^[52]^ While these studies have advanced understanding of biochemical and biophysical signaling between muscles and tendons, no studies to date, to our knowledge, have investigated force transmission from contractile muscle through tendons to a rigid skeleton.^[53]^ Replicating this *in vivo* architecture is critical to engineering a biohybrid MTU for applications in robotics, but remains an unaddressed challenge in the field.

We formulated a synthetic tendon based on an established formulation for a biocompatible tough hydrogel, namely an interpenetrating network of poly (vinyl alchohol) and poly (acrylic acid), PVA:PAA. Previous reports have demonstrated that PVA:PAA hydrogels functionalized with N-Hydroxysuccinimide esters are capable of robust adhesion to biological tissues *ex vivo* and *in vivo*,^[54]^ though adhesion to tissue engineered constructs has not previously been investigated, to our knowledge. We bound NHS-functionalized PVA:PAA onto a polyurethane backing (PUD3 or Hydrothane), laser cut the material into a tendon-like shape, and pressed this synthetic hydrogel tendon mimic to both ends of an engineered muscle tissue strip for 1 minute to form robust adhesive bonds, thus generating a biohybrid MTU (Figure 1C). The muscle tissue was fabricated using our previously established protocols for generating contractile 3D skeletal muscle from C2C12 mouse myoblasts.^[55]^ Briefly, myoblasts were embedded in a fibrin/Matrigel hydrogel, maintained in growth media containing fetal bovine serum for ∼2 days, and differentiated in media containing horse serum and human insulin-like growth factor 1 for ∼10 days (Figure S1). In this study, muscles were fabricated from an established optogenetic C2C12 line that expressed the light-sensitive calcium ion channel ChR2[H134R], enabling non-invasive control of muscle contraction via stimulation with 470 nm blue light.^[5]^.

A quantitative viability assay validated that co-culturing muscle cells with NHS-functionalized PVA:PAA hydrogels did not have a significant impact on cell viability as compared to control muscle cells (Figure 1D), validating the biocompatibility of proposed tendon material. Likewise, tensile test of biohybrid MTUs (n = 6) demonstrated that a 1*x*1mm muscle-tendon interface could withstand forces > 5 mN without delamination, surpassing the magnitude of the typical passive tension and active contraction forces (< 1 mN) generated by mm-scale C2C12-derived muscle^[4,5]^ (Figure 1E) and validating the structural stability of the MTU for applications in robotics.

### 2.2 Mathematically modelling and characterizing biohybrid MTUs

Studying the impact of various MTU design metrics on performance requires accurately assessing how these parameters affect the biohybrid actuator’s contractile magnitude and dynamics. To perform this characterization, we mounted MTUs onto a flexure-based skeleton designed to provide minimal resistance to movement in the direction aligned with muscle contraction, while preventing displacement or rotation along other axes, thus mimicking an articulating joint (**Figure 2**A-B). The skeleton was machined from low-density polyethylene with two rigid pins serving as anchor points for the MTU, where one pin is fixed, and the other pin is attached to the flexure and thus free to move. *In vivo,* MTUs are always held in a state of passive tension to ensure both force generation and force transmission occur along the same axis. We thus integrated an adjustable cam with our flexure-based skeleton to enable pre-tensioning biohybrid MTUs to different degrees.

**Figure 2.**
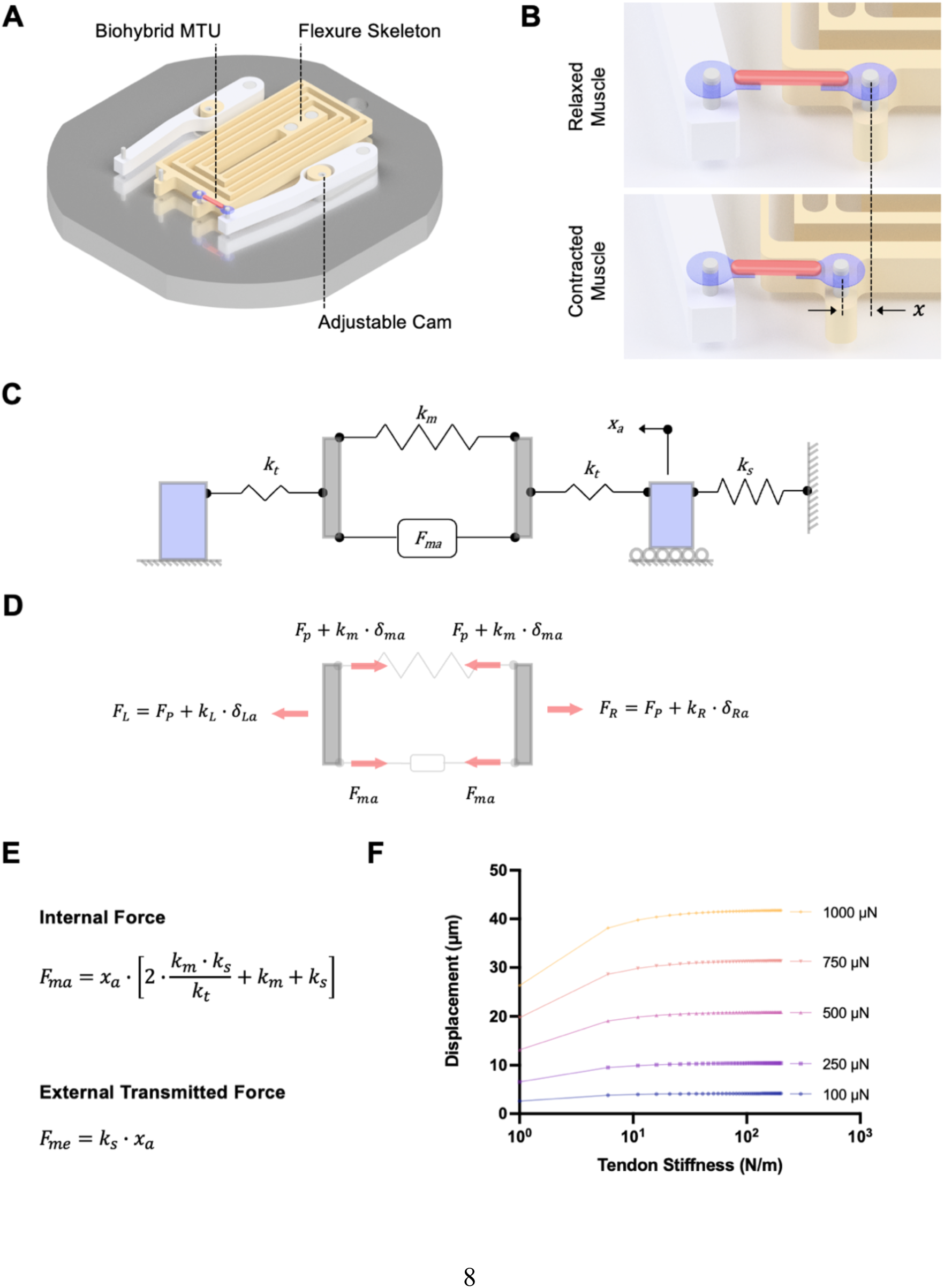
Characterization and mathematical modeling of biohybrid MTU. (**A**) Schematic of flexure skeleton (yellow) mounted on a rigid base (gray). Biohybrid MTUs composed of a muscle strip (pink) bound to a tendon hydrogel (blue) at each end are mounted onto the pins of the flexure skeleton. (**B**) Stimulated muscle contraction displaces the mobile pin of the flexure, enabling accurate visual measurements of muscle contractile deformation and dynamics. (**C**) Mathematical model of biohybrid MTUs and skeletons as elastic springs in series. The tendon is represented by a spring of stiffness *k_t_*, the skeleton is represented by a spring of stiffness *k_s_*, and the muscle (which generates force *F_ma_*) is represented by a spring of stiffness *k_m_*. (**D**) Free body diagram of left and right sides of MTU. (**E**) Equations for internal muscle actuation force and external transmitted force to skeletons. (**F**) Plotting displacement of flexure skeletons, *x_a_*, as a function of tendon stiffness for different internal muscle actuation forces showing that displacement is maximized at tendon stiffness > 100 N/m in our system (*k_m_* = 24 N/m, *k_s_* = 0.3 N/m).

We developed a mathematical model of our system to convert measured displacements of the skeleton’s mobile pin into values of muscle force by modeling the skeleton, tendon, and muscle as elastic mechanical elements in series with spring constants *k_s_*, *k_t_*, and *k_m_* respectively (Figure 2C). Pre-tensioning the MTU generates a preload force, *F_p_*, and stimulating the muscle generates an actuation force, *F_ma_*. The contractile displacement of the muscle, *δ_ma_*, and stretches external elements (i.e. tendon, skeleton) to the left (*δ_La_*) and to the right (*δ_Ra_*) of the muscle. A free body diagram of the system (Figure 2D) combined with compatibility equations (*δ_La_* + *δ_ma_* + *δ_Ra_* = 0) can be combined to obtain equations for the muscle internal actuation force, *F_ma_*, and the muscle external transmitted force, *F_me_*, as a function of the measured displacement of the skeleton’s mobile pin, *x_a_*, as well as *k_s_*, *k_t_*, and *k_m_* (Figure S2, Figure 2E).

Given that the muscle stiffness, *k_m_* (a function of muscle modulus, cross-sectional area, and length), and actuation force, *F_ma_*, are determined by the composition of the tissue (i.e. cell line, extracellular matrix-mimicking hydrogel), these were considered fixed parameters in our model. To optimize tendon design parameters, we plotted skeleton displacement, *x_a_*, as a function of tendon stiffness, *k_t_*, for a range of *F_ma_* from 100-1000 µN (Figure 2F). These models predicted that, for our muscle tissues with *k_m_* ∼24 N/m, tendon stiffnesses *k_t_* > 100 N/m would be expected to maximize displacement of flexure skeletons, *x_a_*, and thus transmitted force, *F_me_* = *k_s_ x_a_*. We conducted a tensile test of PVA:PAA hydrogels with both PUD3 and Hydrothane backings to determine their modulus (2.5 MPa and 6.3 MPa respectively, Figure S3), enabling us to choose tendon geometric parameters that yielded stiffnesses greater than 100 N/m. Specifically, tendons of dimensions 2 *x* 3 *x* 0.3 mm (corresponding to *k_t_* = 500 N/m for PUD3-backed hydrogels and *k_t_* = 1260 N/m for Hydrothane-backed hydrogels) were used in subsequent experiments to characterize the force transmitted from biohybrid MTUs to both compliant and stiff flexure skeletons.

### 2.3 Tuning MTU stiffness and pre-stretch to modulate contractile displacement across frequencies

MTUs *in vivo* are always held in a state of passive tension, motivating us to investigate the impact of pre-stretch on contractile displacement and dynamics. We have previously demonstrated that muscles coupled to compliant flexure-based skeletons (0.3 N/m mechanism stiffness) generate contractile strains that match the strain of free untethered muscle tissues floating in media (i.e. the theoretical maximum of movement) and thus enable characterizing the muscle’s full dynamic range of motion.^[44]^ To mimic *in vivo* design, we thus mounted MTUs with PUD3-backed tendons and optogenetic muscles on flexure skeletons with *k_s_* = 0.3 N/m and quantified mobile pin displacement in response to 1 Hz stimulation as a function of pre-stretch (**Figure 3**A-B).

**Figure 3.**
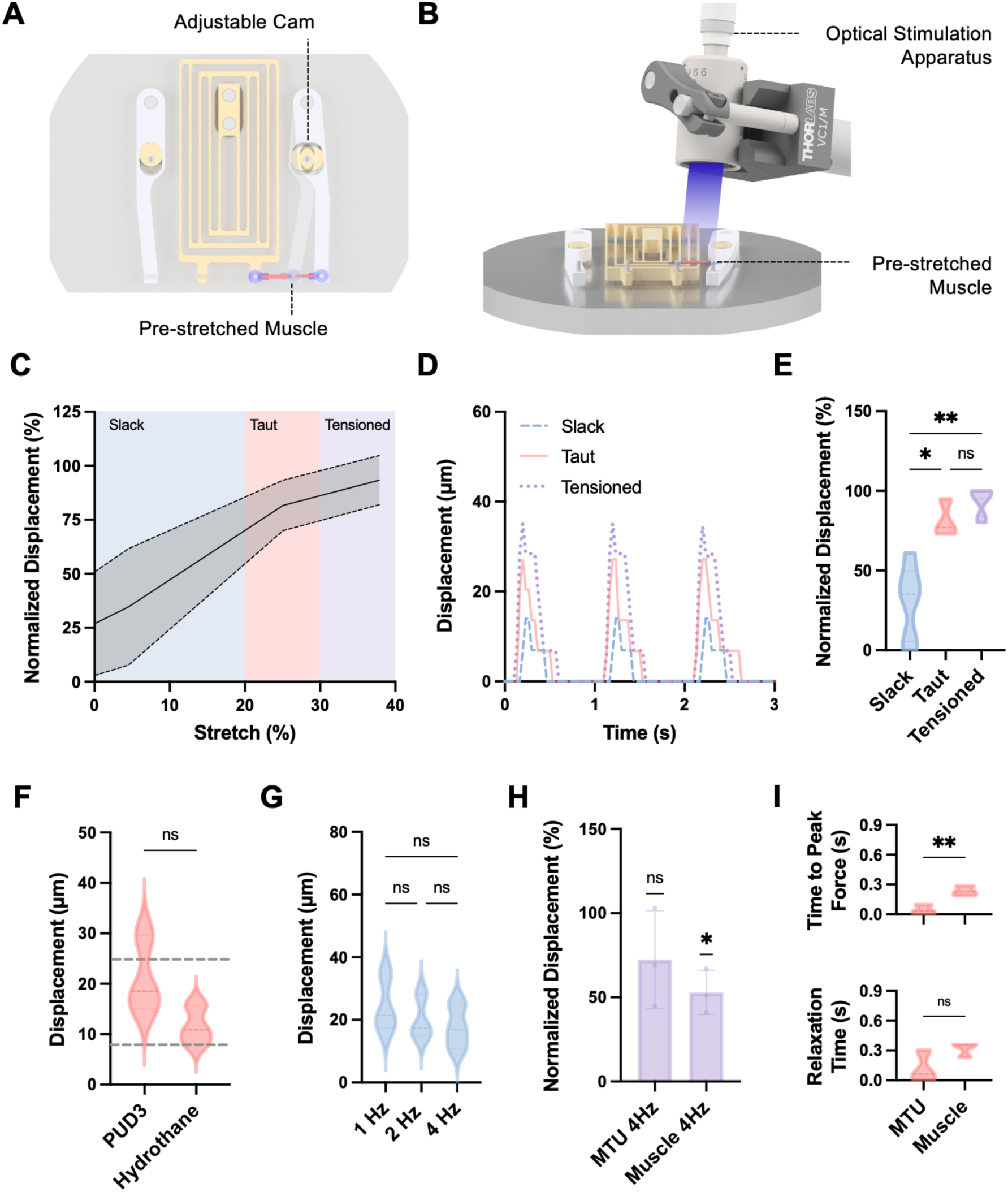
Impact of biohybrid MTU pre-stretch on performance. (**A**) Adjustable cam enables pre-stretching the MTU to different degrees. (**B**) Schematic of apparatus used to stimulate optogenetic muscles within MTUs with 470 nm blue light. (**C**) Average displacement of PUD3-backed MTUs at 1 Hz stimulation normalized to the maximum observed displacement for slack (blue), taut (pink), and tensioned (purple) conditions. Dashed lines represent standard deviation, n = 3 samples. (**D**) Displacement of a representative PUD3-backed MTU in slack, taut, and tensioned conditions in response to 1 Hz optical stimulation. (**E**) Pre-tension significantly impacts PUD-3 backed MTU normalized displacement in response to 1 Hz stimulation. 2-way ANOVA, n = 3, *p < 0.05, **p < 0.01. (**F**) Displacement of taut MTUs with PUD3-backed (500 N/m) and Hydrothane-backed (1260 N/m) tendons in response to 1 Hz stimulation. Dashed lines represent displacement values predicted by our mathematical model for *k_s_* = 0.3 N/m and *F_ma_* ranging from a minimum of 200 µN to a maximum of 600 µN. Welch’s t test, n = 3 per condition. (**G**) Displacement of taut PUD3-backed MTUs in response to 1, 2, and 4 Hz optical stimulation. 2-way ANOVA, n = 3. (**H**) Displacement of taut PUD3-backed MTUs on skeletons compared to muscles placed directly on skeletons in response to 4 Hz stimulation, normalized to displacement at 1 Hz stimulation. One sample t test compared to theoretical mean of 100%, *p < 0.05. (**I**) Time-to-peak-force (top) and relaxation time to baseline (bottom) for taut PUD3-backed MTUs versus muscles alone at 1 Hz stimulation. Welch’s t test, n = 3, **p < 0.01.

MTUs were stretched from 0-45% of their original length, and skeleton displacements were measured for slack (defined as 0-20% stretch), taut (20-30% stretch), and tensioned (30-45% stretch) conditions. We observed that MTU contractile displacement increased significantly between slack MTUs and taut MTUs, but that tensioning MTUs further did not significantly enhance performance (Figure 3C-E, Video S1). Notably, taut MTUs with both PUD3-backed and Hydrothane-backed tendons generated contractile displacements that were not significantly different from each other (Figure 3F). Videos of free-floating muscle tissues (n = 6) in response to 1 Hz stimulation were used to empirically determine a range of *F_ma_* for our tissues of 400 ± 200 µN. The median skeleton displacements in both PUD3-backed and Hydrothane-backed MTU groups fell within the range of displacement predicted by our mathematical model for these *F_ma_* values (dashed lines in Figure 3F). Together, these results indicated that our mathematical model could predict skeleton displacement given known *k_s_*, *k_t_*, *k_m_*, and *F_ma_*. Moreover, they indicated that increasing tendon stiffness further was unlikely to increase observed displacement. Taut MTUs with PUD3-backed tendons were thus used for all following experiments.

To further characterize the impact of stimulation frequency on MTU contractility, we measured and compared skeleton displacement in response to higher frequency stimulation (Figure 3G, Video S2). Interestingly, we observed that MTUs demonstrated similar contractile displacement at 4 Hz as compared to 1 Hz, in contrast to muscles placed directly on skeletons which demonstrated a smaller dynamic range of motion at higher frequencies (Figure 3H). We attributed these differences to decreases in the time-to-peak-force of MTUs as compared to muscles (Figure 3I), indicating that the addition of synthetic tendons modulates the bioactuator’s contractile dynamics.

### 2.4 Determine fatigue characteristics of biohybrid MTUs

Practical use of MTUs in a biohybrid robot requires establishing whether the muscle-tendon-skeleton interface withstands repeated loads over many cycles of contraction, and whether the muscle itself demonstrates fatigue over long periods of continuous use. To investigate these phenomena, we conducted a fatigue test of MTUs with tendons in a taut configuration by subjecting them to continuous stimulation at 2 Hz or 4 Hz for 30 minutes, corresponding to 3600 and 7200 loading cycles respectively (**Figure 4**A).

**Figure 4.**
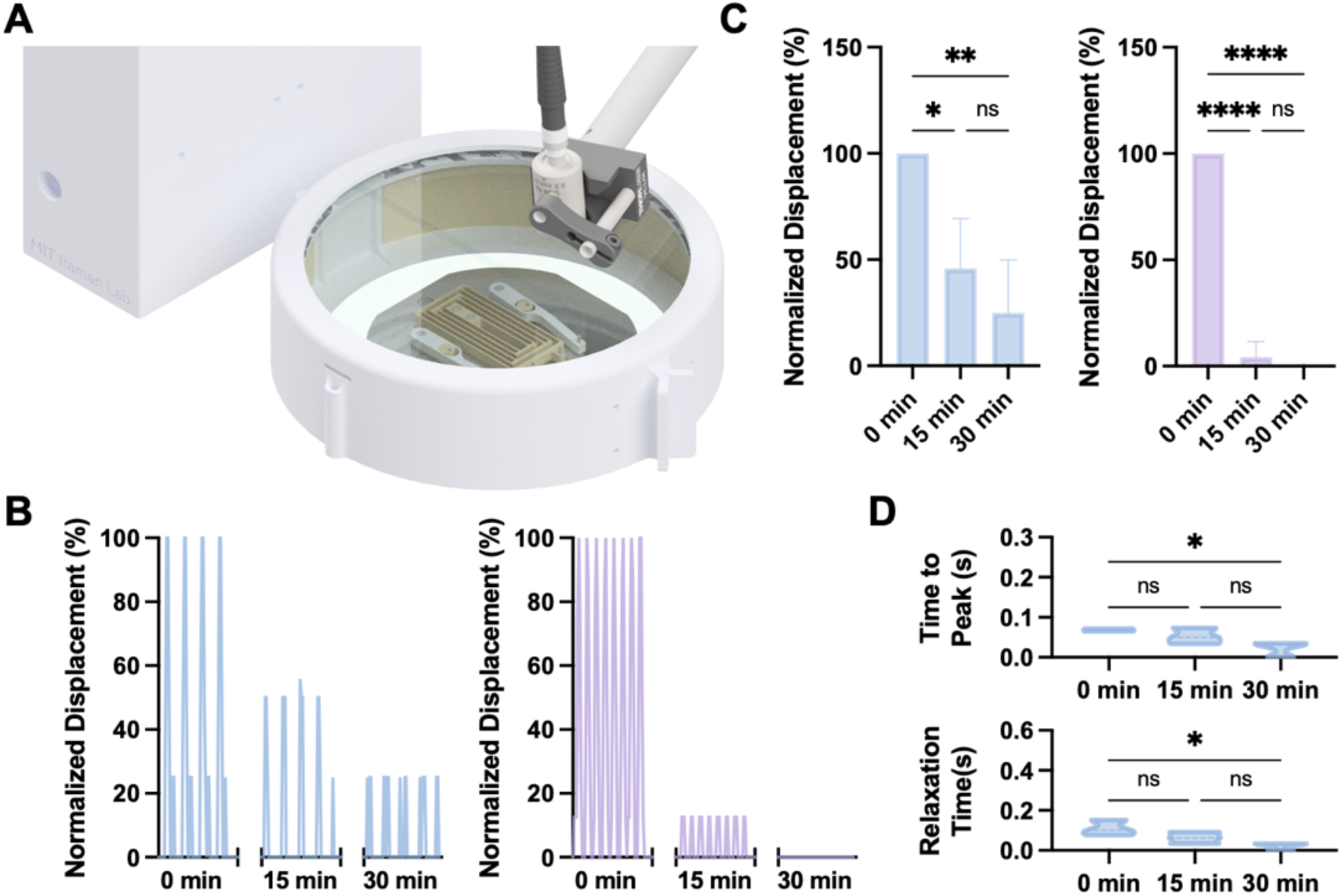
Characterization of biohybrid MTU fatigue. (**A**) Schematic of custom incubation apparatus used to perform temperature-controlled fatigue testing of MTUs. (**B**) Representative traces of contractile displacement, normalized to t = 0 values for 2 Hz (left) and 4 Hz (right) continuous stimulation at 0, 15, and 30 minutes (∼2 seconds shown per time point). (**C**) Contractile displacement for 2 Hz (left) and 4 Hz (right) continuous stimulation normalized to displacement at t = 0 minutes. 2-way ANOVA, n = 3, *p < 0.05, **p < 0.01, ****p < 0.0001. (**D**) Time-to-peak-force (top) and relaxation time (bottom) for MTUs at 2 Hz stimulation. 2-way ANOVA, n = 3, *p < 0.05.

Monitoring muscle displacement at 0, 15, and 30 minutes revealed that, in both stimulation conditions, the muscle-tendon interface demonstrated no signs of delamination (Video S3). However, MTU contractile displacement dropped significantly over the 30-minute observation period in both stimulation conditions (Figure 4B-C, Video S3). Contractile dynamics of MTUs were measured at 0, 15, and 30 minutes for the 2 Hz condition, as it generated measurable deformation throughout the testing period. Both time-to-peak-force and relaxation time decreased significantly over 30 minutes (Figure 4D), though this could be attributed to the reduced displacement per stroke. Notably, in our previous studies of muscle endurance, we observed that muscle actuators placed directly on flexure skeletons demonstrated significant fatigue (> 80% drop in force) in response to 4 Hz continuous stimulation for 30 minutes.^[44]^ This similarity between muscle fatigue data and MTU fatigue data indicates that the actuator’s fatigue behavior is likely tied to the endurance of the muscle (typically determined by the availability of nutrients and distribution of muscle fiber types within the tissue^[56,57]^), rather than fatigue-induced changes to the muscle-tendon interface.

### 2.5 Deploying MTUs on a high-stiffness skeleton to increase transmitted force

Prior studies of biohybrid robots couple muscles directly to skeletons. As a result, these studies deploy muscles on low-stiffness mechanisms (characterized by large displacements and low transmitted force), in an effort to prevent stress concentrations at the muscle-skeleton interface that could damage tissues. However, plotting the ratio of the muscle’s transmitted force, *F_me_*, to the internal actuation force, *F_ma_*, as a function of *k_s_* shows that, in our system, transmitted force increases with increasing skeleton stiffness up to a plateau of ∼85% (**Figure 5**A). Maximizing the force exerted by the muscle on the skeleton thus requires significantly increasing skeleton stiffness, while still preserving tissue integrity and the ability to resolve contractile displacement.

**Figure 5.**
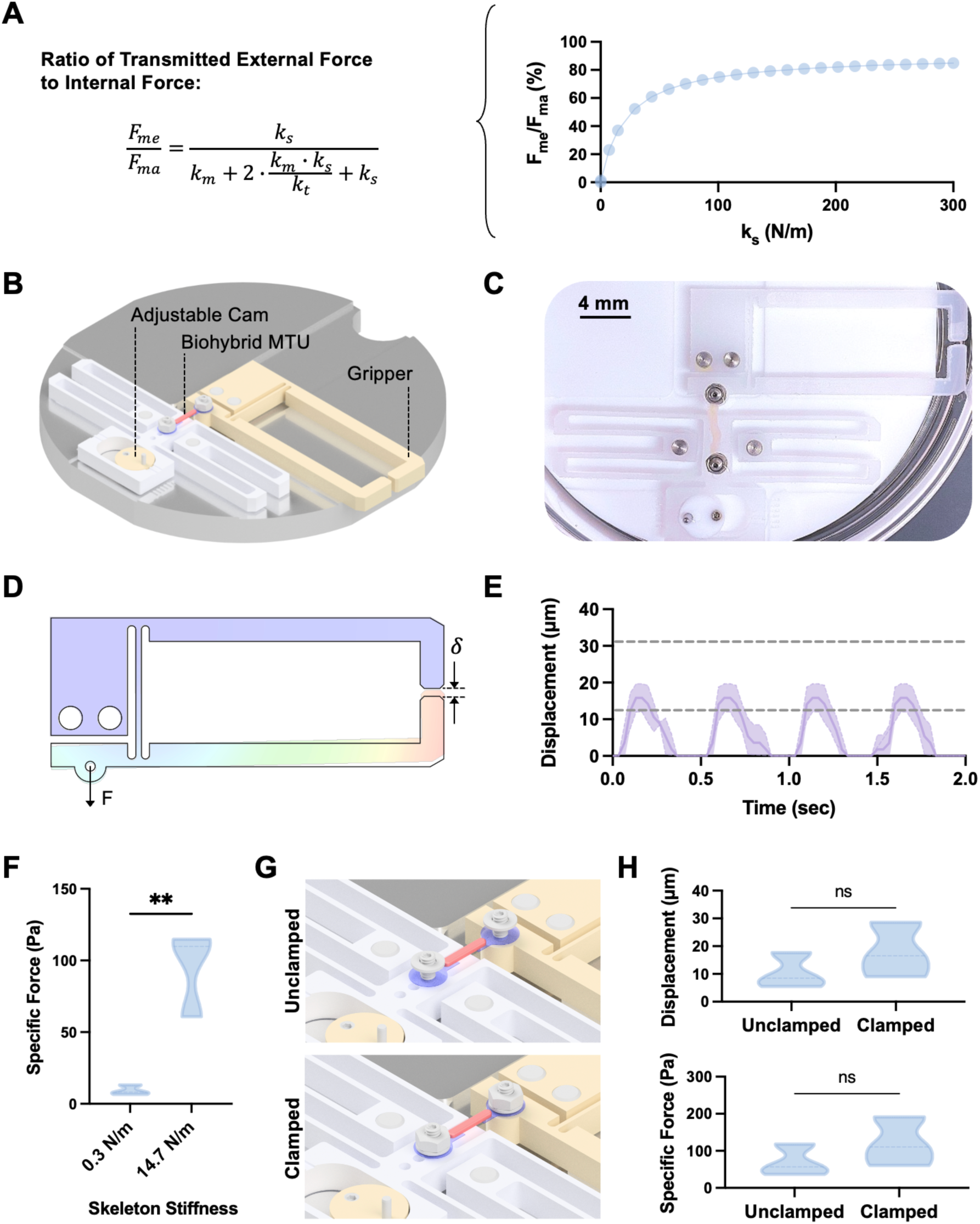
Characterization of biohybrid MTU fatigue. (**A**) Left: Equation showing ratio of MTU transmitted force, *F_me_*, to internal actuation force, *F_ma_*. Right: Plot of *F_me_/F_ma_* as a function of skeleton stiffness, *k_s_*. (**B**) Schematic of high-stiffness gripper design with integrated cam to enable MTU pre-tensioning. (**C**) Real image of biohybrid MTU mounted onto gripper. (**D**) Finite element analysis simulation showing how muscle contraction force, *F*, creates deflection, *δ*, at the tip of the gripper. (**E**) Displacement of gripper tip in response to optical stimulation of biohybrid MTUs, with shaded area showing range of movement for n = 3 samples. Dashed lines represent displacement values predicted by our mathematical model for *k_s_* = 14.7 N/m and *F_ma_* ranging from a minimum of 200 µN to a maximum of 600 µN. (**F**) Specific force generated by biohybrid MTUs on high-stiffness gripper skeleton as compared to low-stiffness skeleton. Unpaired t-test, n = 3, **p < 0.01. (**G**) Schematic of clamping approach used to prevent rotating or slipping about the tendon/skeleton interface in the gripper. (**H**) Clamping the MTU on the gripper does not significantly increase displacement (top) or specific force (bottom), as compared to unclamped samples. Unpaired t-test, n = 3.

We designed a flexure-based skeleton with a stiffness of 14.7 N/m, ∼50X stiffer than our previous skeleton design, in the form of a two-armed gripper (Figure 5B-C). Given that increasing skeleton stiffness was also expected to reduce contractile displacement, our stiff skeleton design integrated long gripper arms that amplified muscle contractile displacement by ∼2.8X to ensure accurate visual measurements (Figure 5D). The stiff gripper skeleton also integrated an adjustable cam to enable pre-stretching MTUs.

We measured the displacements of the gripper arm in response to MTU stimulation (Figure 5E, Video S4) and observed that empirically measured median displacements fell within the range predicted by our mathematical model for *F_ma_* = 400 ± 200 µN. Also as predicted by our model, MTUs transmitted significantly higher specific forces (contractile force normalized by muscle cross-sectional areas) to our high-stiffness skeletons as compared to our previous low-stiffness skeleton design (Figure 5F). Notably, MTUs on high-stiffness skeletons demonstrated an *F_me_*/*F_ma_* of ∼37%, as compared to ∼1.3% on low-stiffness skeletons, corresponding to a ∼29X increase in force transmission. Changing skeleton stiffness did not impact MTU contractile dynamics (Figure S4A).

To examine whether tendons were slipping or rotating against the interface with the high-stiffness skeleton, we developed a method of mechanically clamping the tendons and skeletons to each other with a flat washer and nut (Figure 5G). Interestingly, we observed that clamping the tendon-skeleton interface did not significantly increase the displacement of the gripper arms or the specific force transmitted from the MTU to the skeleton (Figure 5H), indicating minimal slipping at the tendon-skeleton interface in our system, possibly due to the induced pre-stretch in the MTU. Furthermore, clamping did not change the contractile dynamics (Figure S4B).

## 3.0 Discussion

Biohybrid robots powered by skeletal muscle typically rely on strip- or ring-based designs that couple these actuators directly to skeletons. This architecture wastes a large percentage of the muscle tissue by using it to perform a non-contractile structural role, and also constrains the upper limit of skeleton stiffness to prevent stress concentrations at the muscle-skeleton interface. In this study, we take inspiration from the native muscle-tendon unit, or MTU, by fabricating a biohybrid MTU composed of a linear soft muscle actuator with a tough hydrogel tendon at each end. Since the PVA:PAA hydrogel tendons have much higher fracture strengths than the tissues themselves,^[54,58]^ and can form robust adhesive bonds with the tissues (Figure S1), the MTU can withstand high stress concentrations at the MTU-skeleton interface, thus enabling deployment on high-stiffness mechanisms. Moreover, using tendons to perform a structural role ensures that nearly all the muscle tissue is used to generate useful actuation, thus limiting tissue waste.

We measured the impact of various design parameters on actuator performance. Our experiments corroborated our mathematical model by showing that effective MTU function requires tendons that are much stiffer than muscle. Furthermore, we show that pre-tensioning actuators to a “taut” configuration (corresponding to ∼20% stretch as compared to the passive length) maximizes displacement. Interestingly, we observed that MTUs had lower time-to-peak force than muscles alone, and thus demonstrated larger displacements at high-frequency stimulation. However, similar to muscles alone, MTUs displayed significant fatigue over 30 minutes of continuous high-frequency stimulation. To showcase the modularity of our MTUs, we deployed them on two skeleton designs (with ∼50X difference in stiffness), and validated our mathematical model of muscle-tendon-skeleton interactions to show that the force transmitted from muscles to skeletons increases with increasing mechanism stiffness.

To highlight the efficiency advantage of MTUs over traditional methods of direct muscle-skeleton coupling, we compared the power-to-weight ratio of our actuator to previous actuators from ourselves and other researchers using the same muscle formulation (C2C12 myoblasts in a fibrin/Matrigel hydrogel) (**Figure 6**). This calculation highlighted that MTUs outperform previous designs that rely on direct muscle-skeleton coupling by ∼11X on average, likely because they limit wasteful use of muscle tissue to perform structural roles.

**Figure 6.**
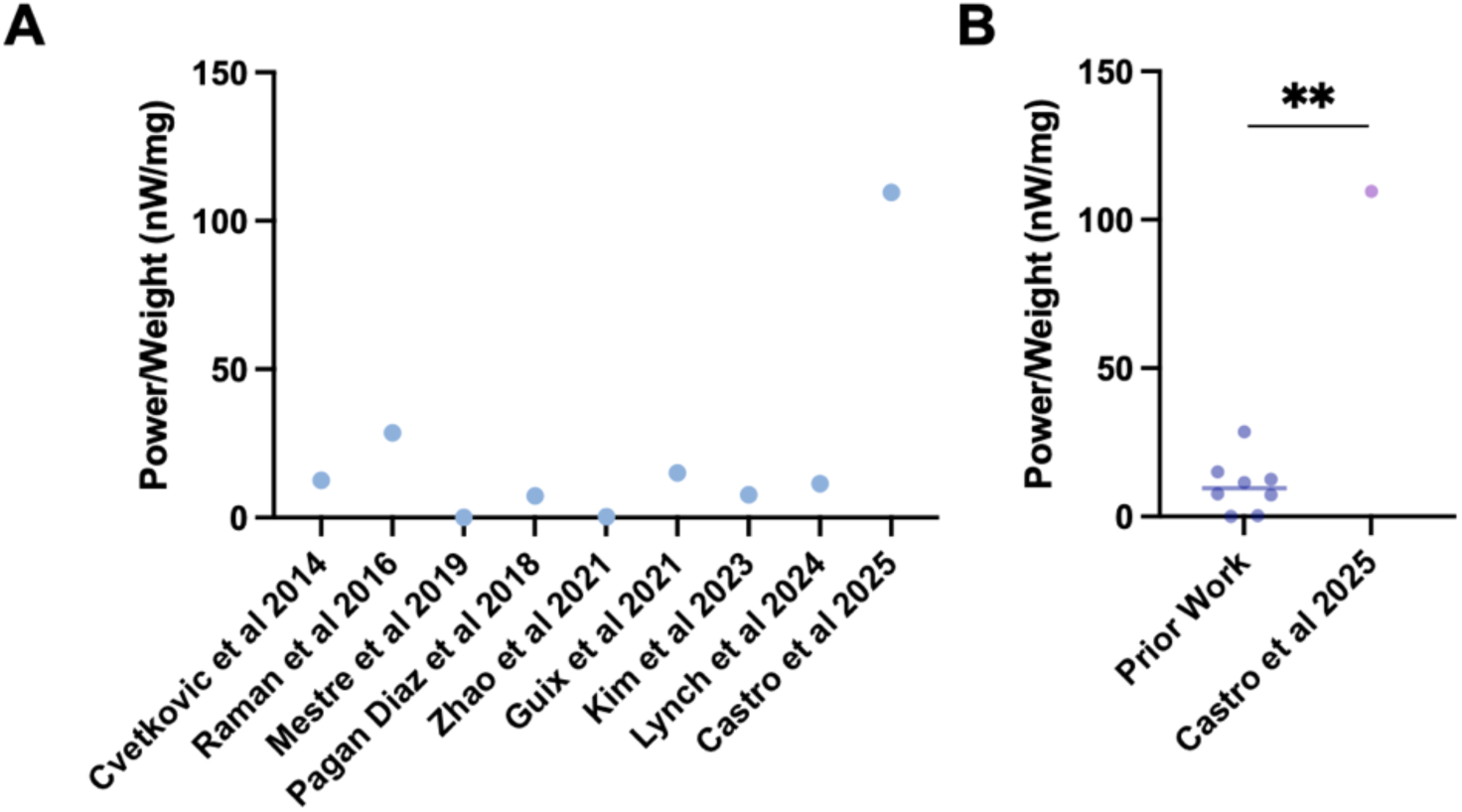
Power-to-weight ratio of biohybrid MTUs. (**A**) Comparison of biohybrid MTU (Castro et al 2025) power-to-weight ratio as compared to prior demonstrations of muscles placed directly on skeletons using the same muscle formulation (C2C12 myoblasts in fibrin/Matrigel hydrogel).^[4–7,29,31,44,59]^ (**B**) Biohybrid MTUs significantly outperform prior work in power-to-weight ratio. One sample t and Wilcoxon test comparing median of prior work to current work, 109.6 nW/mg.

A limitation of our study is that we have not fully replicated *in vivo* MTU design, in which the myotendinous junction has a stiffness gradient from muscles to tendons, rather than a discrete jump in stiffness as in our design. It is possible that future longitudinal studies of our MTUs will show that a sudden jump in stiffness at the muscle-tendon interface harms muscle fiber maturity and strength, motivating the development of PVA:PAA tendon hydrogels with spatially varying stiffness. An additional limitation of our study is that we did not explore higher stiffness skeletons (> 100 N/m) that could maximize the ratio of *F_me_*/*F_ma_*, given the difficulty of visually measuring displacement in these configurations. Future studies that leverage strain sensors to perform more automated and accurate measurements of muscle displacement would enable exploring the MTU’s contractile behavior on even higher-stiffness mechanisms.^[31,32]^ Finally, we use an immortalized mouse cell line to fabricate our tissues, but other studies have shown higher contractile deformations generated by primary cells^[22]^ and human cells.^[24]^ Future studies that use higher-performing cell sources may thus drive a deeper understanding of the upper limits of MTU function.

The modular design of our biohybrid MTU ensures that muscle actuators do not need to be redesigned for each end-use application, but rather can be readily coupled to a variety of skeleton designs integrating mobile pins to enable different functional behaviors, as outlined in Figure 3 and Figure 5. This mimics the current standard integration architecture for other non-biological actuators, such as pneumatic systems, which can be “plugged in” to different robot architectures without requiring actuator re-design. In future, this modularity and design standardization can be leveraged by multiplexing arrays of MTUs to power multi degree-of-freedom motion in more complex robotic mechanisms. We anticipate that biohybrid muscle-tendon units will enhance the modularity, efficiency, and multifunctionality of muscle-powered robots in the coming years.

## 4.0 Materials and Methods

### Fabrication of Synthetic Tendon

Synthetic tendons were fabricated from a hydrogel composed of interpenetrating networks of poly(vinyl alcohol) (PVA) and poly(acrylic acid) (PAA) functionalized for tissue adhesion with N-Hydroxysuccinimide (NHS) esters. Using previously reported protocols ^[1,2]^, the adhesive layer is prepared by mixing 35% (w/w) acrylic acid (Sigma-Aldrich) with 7% (w/w) PVA (Mw = 146,000–186,000, 99+% hydrolysed, Sigma-Aldrich) in deionized water, along with additives of 0.2% (w/w) α-ketoglutaric acid (Sigma-Aldrich) and 0.05% (w/w) *N,N’*-methylenebisacrylamide (MBAA) (Sigma-Aldrich). Then, 30mg of *N*-hydroxysuccinimide ester functionalized acrylic acid (Thermo Scientific) is mixed to 1ml of the above solution, poured onto glass mold plates with 100µm spacers, and cured under UV for 30 min (354nm wavelength, 12W). After curing, a polyurethane backing of PUD3 (Hydromed D3) solution is spin coated onto the hydrogel at 200rpm for 60 seconds. The Hydromed D3 solution is composed of 10% (w/w) Hydromed D3 (AdvanSource Biomaterials) in reagent alcohol (90% ethanol, 5% isopropanol, 5% methanol) (BDH Chemicals). The hydrothane solution is composed of 20% Hydrothane AL 25-80A (AdvanSource Biomaterials) in 1:1 tetrahydrofuran (THF) (TCI) and *N,N*-Dimethylformamide (DMF) (Sigma-Aldrich). Once solidified, the synthetic tendon is prestretched in length and width equal to the equilibrium swelling ratio through manual and automated stretching setups. After drying under airflow and in a vacuum desiccator, the synthetic tendon can be laser cut into desired shapes and stored at - 20C and vacuum sealed until use.

### Fabrication of Flexure Skeletons

The flexure component of both compliant and stiff gripper skeletons was manufactured from low-density polyethylene (LDPE). Prior to machining via precision CNC micro-milling, the LDPE was stress relieved to remove residual stresses. For the compliant skeleton, the conjugated beams of the flexure were machined to a thickness of 0.5mm and beam lengths along the conjugated chain varied between 25mm and 40mm. All beams were machined to a height of 3.35mm. For the stiff gripper skeleton, the arm beam was machined to be 0.71 mm wide and 17 millimeters long. Cavities were machined in the underside of the gripper arms to enable the capture of air bubbles whose volume was calculated to make the arms neutrally buoyant within the fluid cell culture medium. All beams were machined to a height of 3.35mm. To prevent the flexure beams from deflecting during machining, the LDPE was fixed in place via double sided tape and the machining process was optimized to minimize cutting forces that would deflect or damage the flexure beams. A light finishing pass was conducted on all beam side edges to prevent significant ‘spring back’ due to deformation of the beams under cutting forces. Both compliant and stiff flexures were mounted onto a base (aluminum for compliant flexure, Teflon for stiff flexure) alongside adjustable cam arms for machined from Teflon. Prior to use in experiments with MTUs, the adjustable flexure skeleton was washed in 1x DPBS, submerged in 70% ethanol and exposed to UV light for 1 hour, and then washed again with 1x DPBS.

### Fabrication of Muscles

Skeletal muscle rings were fabricated by following a previously published protocol ^[5,55]^. C2C12 myoblasts genetically engineered to express an optogenetic calcium ion channel, ChR2[H134R], were cultured in growth medium composed of DMEM with additions of 4.5g/L of glucose, L-glutamine, and sodium pyruvate (Corning) supplemented with 10% (v/v) FBS (Life Technologies), 1% (v/v) penicillin/streptomycin (Fisher Scientific), and 1% (v/v) L-glutamine (Fisher Scientific). Upon reaching at least 70% confluency, cells were trypsinized for tissue fabrication. At least one day prior to muscle ring fabrication, high concentration Matrigel (Matrigel Basement Membrane Matrix, Corning) was thawed at 4°C. In addition, fibrinogen (Sigma Aldrich) was weighed and stored in a conical at -20°C. On the day of muscle ring fabrication, thrombin (Sigma Aldrich) stock solution (100U/mL in 0.1% wt/v bovine serum albumin (BSA), Sigma Aldrich), and supplemented growth media containing aminocaproic acid (6-Aminocaproic acid, Sigma Aldrich) stock solution (50mg/mL in DI) for a final concentration of 1mg/mL were thawed and kept in an ice bath. Sterilized molds made of poly (dimethyl siloxane) (PDMS) were placed in 6-well plates and submerged in 8 mL of 1% w/v BSA in 1x DPBS for at least 1 hour. Cultured cells were trypsinized (TrypLE Express, Gibco), harvested and counted into aliquots of 3 *x* 10^6^ cells per vial. Fibrinogen was dissolved in supplemented growth media at a concentration of 8mg/mL and vortexed thoroughly, then set on ice. Cell aliquots were centrifuged at 1000 rpm for 5 minutes at room temperature to pellet the cells. The supernatants of these aliquots were aspirated off, and the cell pellets were resuspended in 59 µl of the supplemented growth media. Matrigel at 4°C was then also set on ice. Once the cells were resuspended, per aliquot prepared, 1µL of the thrombin stock solution, 90µL of Matrigel and 150µL of the fresh fibrinogen stock solution were added to the cell suspension in order. The cell/hydrogel suspension was used to fill the muscle molds and then incubated at 37°C with 5% CO2 for 1 hour and then submerged in growth medium. Tissues were maintained on PDMS molds in growth medium for 2 days and then cultured in differentiation medium composed of DMEM with 4.5g/L of glucose, L-glutamine, and sodium pyruvate, supplemented with 10% (v/v) Horse Serum (Gibco), 1% (v/v) penicillin/streptomycin, 1% (v/v) L-glutamine, 1mg/mL of aminocaproic acid and 5ng/mL of IGF-1 (Sigma Aldrich).

### Fabrication and Stimulation of Muscle Tendon Units (MTUs)

MTUs were fabricated on the day they were used. First, synthetic tendon stock was retrieved from -20°C immediately before fabrication. A muscle tissue was removed from its PDMS mold and placed in an empty petri dish. Tendon strips were placed with tweezers on the muscle strip so that the adhesive side contacted the muscle, with the polyurethane backing on the opposite side. Gentle pressure was then applied at each end interface with tweezers for at least 1 min to assist with adhesion. Once each end had been bound with tendons, the completed unit was then mounted onto flexures, pre-stretched to the desired degree via the adjustable cam, and stimulated with a collimated 470-nm fiber optic LED (ThorLabs), with 1, 2, or 4 Hz square waves with a 20% duty cycle.

### Tendon Adhesion and Modulus Testing

For adhesion testing, two T-brackets from the tensile tester were removed from the setup, cleaned, and sections of double-sided scotch tape were added to the center of each testing plate. The tendons in MTUs were adhered to the T-brackets via the double-sided scotch tape. The assembly was then inserted into an Instron tensile tester and stretched at a rate of 0.01mm/s. For modulus testing, tendons were fabricated in a dog-bone format (1.5cm wide and 5cm long), allowed to swell in media for 24 hours prior to testing, and then mounted into an Instron tensile tester and stretched at a rate of 1.3 mm/s. The resulting force-displacement curve was recorded. Unstretched sections of each sample were cut and fixed between glass slides and thickness was measured under a microscope to calculate cross-sectional area and convert force to stress. A Neo Hookean model was used to convert measured stress and stretch into values of elastic modulus.

### Viability Assay

C2C12 myoblasts were cultured in a 24-well plate, with each well cultured at the same cellular density. Synthetic tendons were into these wells for 24 hours. A colorimetric MTS cell viability assay (CellTiter 96, Promega) was performed on the wells (n=3 for both tendon-exposed and control). After incubation with the MTS compound, media samples were extracted from each well and placed in a 96-well plate for absorbance analysis.

### Fabrication of Incubation Chamber for Fatigue Testing

The custom incubator for long-term MTU use includes an electronics housing for control circuitry and a circular chamber with a removable lid containing heating and sensing elements. Both the chamber and housing were 3D printed using resin on a Stratasys J35 Pro Polyjet printer (White, VeroUltra), while the lid was laser-cut from 0.125” clear acrylic (McMaster-Carr). To improve heat distribution from the heaters, the internal walls of the chamber were lined with 0.005” copper strips. Two flexible resistive heaters with integrated 10 kΩ NTC thermistors (HT10K, Thorlabs) were adhered to the copper lining. An additional 10 kΩ NTC thermistor (372, Adafruit) was used to monitor the internal environmental temperature. A control circuit was designed to modulate power to the heaters from a 24V DC power supply, governed by a PWM voltage from an Arduino Nano. The Arduino calculates the thermistor resistances, from which temperature is derived using the Steinhart-Hart equation. Two PID controllers run simultaneously, one limiting the maximum power to the heaters based on heater temperatures, while the other requests a PWM duty-cycle to the heaters based on the environmental temperature. The combination of these controllers provides both safety against overheating and a fast rise time, while maintaining long-term temperature precision and stability.

### MTU Clamping

The sterilized gripper skeleton was placed in a dish and submerged in warm media. MTUs were mounted onto the pins of the gripper and pre-stretched with the adjustable cam. Washers and threaded nuts were then mounted onto the pins and manually tightened until the nut could no longer rotate. Videos of unclamped and clamped contraction were recorded in response to optical stimulation.

### Data Collection and Analysis

Videos of each experiment were acquired on an iPhone 11, using a Zeiss Stemi 305 Stereoscope with an iDu Optics LabCam attachment in one of its eye pieces. Samples to be recorded were placed under the stereoscope and were illuminated with its equipped overhead light. Recorded videos were converted to .avi file type through FFMPEG processing and analyzed in ImageJ. For displacement tracking, A MATLAB script that utilizes cross-correlation for image analysis was used to track the mobile pin of the compliant flexure or the gripper arm of the stiff flexure. The script tracks the position of the selected feature and records its pixel position in both X and Y dimensions. The resulting arrays of coordinates were then zeroed off the mode of each dimension. Total displacement was then calculated by the RMS of the X and Y dimensions, resulting in a displacement trace, showing the absolute motion of the flexure pin over time due to MTU contraction. Final displacement traces were then compared to the raw X and Y traces and verified for proper tracking. To determine the unique scaling for each video that depended on the microscope objective, LabCam objective, and phone zoom, pixels were converted to µm in each video by taking a known dimension, like the diameter of the pin on the flexure (1000 µm), to set the scale in ImageJ.

### Statistical Analysis

All statistical tests and sample sizes are stated in figure captions.

## Acknowledgements

We would like to acknowledge Dr. Ronald Heisser for technical discussions regarding mechanism design. This work was funded in by a Department of Defense Army Research Office grant W911NF-22-1-0126 P00003 (RR), an MIT Research Committee NEC Corporation Fund Grant (RR), and the National Science Foundation Graduate Research Fellowship Program (MB, SK).

