## Supplemental Information for "Biohybrid tendons enhance the power-to-weight ratio and modularity of muscle-powered robots"

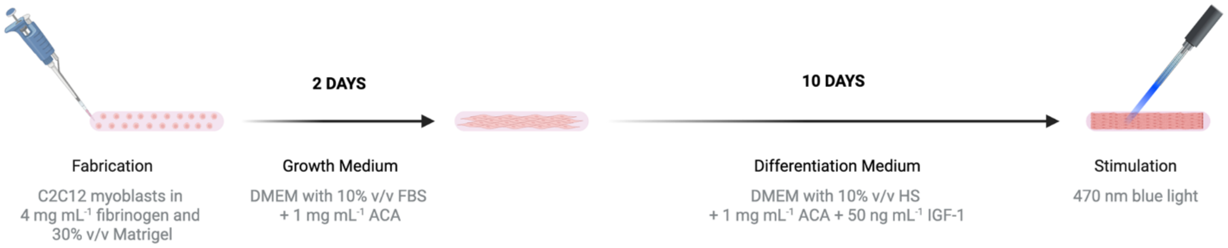


**Figure S1. Schematic of muscle fabrication.** C2C12 myoblasts are embedded in a fibrin/Matrigel hydrogel, maintained in growth medium for 2 days, and differentiated for 10 days prior to MTU fabrication and testing.

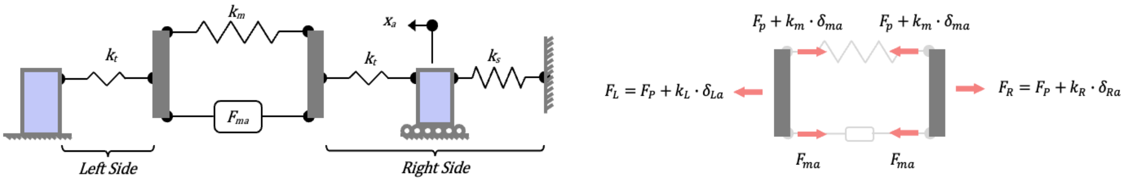


### STIFFNESS DEFINITIONS

*STIFFNESS OF ELEMENTS LEFT OF THE MUSCLE*

$$k_L = k_t$$

*STIFFNESS OF ELEMENTS RIGHT OF THE MUSCLE*

$$k_R = \left(\frac{1}{k_t} + \frac{1}{k_s}\right)^{-1} = \frac{k_t \cdot k_s}{k_t + k_s}$$

$k_t$ = Tendon stiffness
$k_s$ = Skeleton stiffness
$k_L$ = Stiffness of elements left of muscle
$k_R$ = Stiffness of elements right of muscle

### COMPATIBILITY DURING ACTUATION

*ELONGATIONS OF ELEMENTS*

$\delta_{La} + \delta_{ma} + \delta_{Ra} = 0$; Elongations sum to 0 as they're chained between fixed ends

*ELONGATIONS OF ELEMENTS RUGHT OF MUSCLE*

$$\delta_{Ra} = \delta_{ta} + \delta_{sa} = \delta_{ta} + x_a$$

$\delta_{La}$ = Elongation of elements left of muscle
$\delta_{Ra}$ = Elongation of elements right of muscle
$\delta_{ma}$ = Muscle elongation due to actuation
$\delta_{ta}$ = Tendon elongation due to actuation
$\delta_{sa}$ = Skeleton elongation due to actuation
$x_a$ = Pin displacement due to actuation

### STATICS DURING ACTUATION

*SUM OF FORCES ON RIGHT AND LEFT OF MUSCLE WHILE MUSCLE IS ACTUATED*

$$F_L = F_R \rightarrow F_P + k_L \cdot \delta_{La} = F_P + k_R \cdot \delta_{Ra} \rightarrow k_L \cdot \delta_{La} = k_R \cdot \delta_{Ra} \rightarrow \delta_{La} = \delta_{Ra} \cdot \frac{k_R}{k_L}$$

$F_L$ = Force left of muscle
$F_R$ = Force right of muscle
$F_P$ = Preload force

### COMPATIBILITY + STATICS DURING ACTUATION

*ACTUATED ELONGATION OF MUSCLE AND ELEMENTS RIGHT OF MUSCLE*

$$\delta_{La} + \delta_{Ra} = -\delta_{ma} \rightarrow \delta_{Ra} \cdot \frac{k_R}{k_L} + \delta_{Ra} = -\delta_{ma} = \delta_{Ra}\left(\frac{k_R}{k_L} + 1\right) \rightarrow -\delta_{ma} = \delta_{Ra}\left(\frac{k_R + k_L}{k_L}\right)$$

*TENDON ELONGATION AND PIN DISPLACEMENT DURING ACTUATION*

$$F_{ta} = F_{sa} \rightarrow k_t \cdot \delta_{ta} = k_s \cdot \delta_{sa} \rightarrow \delta_{ta} = \delta_{sa} \cdot \frac{k_s}{k_t} \rightarrow \delta_{ta} = x_a \cdot \frac{k_s}{k_t}$$

$F_{ta}$ = Force increment in tendon due to actuation
$F_{sa}$ = Force increment in skeleton due to actuation

### SUM OF FORCES AT RIGHT MUSCLE-TENDON INTERFACE

$$F_P + k_m \cdot \delta_{ma} + F_{ma} = F_P + k_R \cdot \delta_{Ra} \rightarrow F_{ma} = F_P + k_R \cdot \delta_{Ra} - F_P - k_m \cdot \delta_{ma} \rightarrow F_{ma} = k_R \cdot \delta_{Ra} + k_m \cdot (-\delta_{ma})$$

Substitute: $-\delta_{ma} = \delta_{Ra}\left(\frac{k_R + k_L}{k_L}\right)$

$$F_{ma} = k_R \cdot \delta_{Ra} + k_m \cdot \delta_{Ra} \cdot \left(\frac{k_R + k_L}{k_L}\right) = \delta_{Ra} \cdot \left(k_m \cdot \left(\frac{k_R + k_L}{k_L}\right) + k_R\right) = \delta_{Ra} \cdot \left(\frac{k_m}{k_L} \cdot k_R + k_m + k_R\right)$$

Substitute: $\delta_{Ra} = \delta_{ta} + x_a$

$$F_{ma} = (\delta_{ta} + x_a) \cdot \left(\frac{k_m}{k_L} \cdot k_R + k_m + k_R\right)$$

Substitute: $\delta_{ta} = x_a \cdot \frac{k_s}{k_t}$

$$F_{ma} = \left(x_a \cdot \frac{k_s}{k_t} + x_a\right) \cdot \left(\frac{k_m}{k_L} \cdot k_R + k_m + k_R\right) = x_a \cdot \left(\frac{k_s + k_t}{k_t}\right) \cdot \left(\frac{k_m}{k_L} \cdot k_R + k_m + k_R\right)$$

$$F_{ma} = x_a \cdot \left(\frac{k_s + k_t}{k_t}\right) \cdot \left(\frac{k_m}{k_L} \cdot k_R + k_m + k_R\right)$$

Substitute: $k_L = k_t$ Substitute: $k_R = \frac{k_t \cdot k_s}{k_t + k_s}$

$$F_{ma} = x_a \cdot \left(\frac{k_s + k_t}{k_t}\right) \cdot \left(\frac{k_m}{k_t} \cdot \frac{k_t \cdot k_s}{k_t + k_s} + k_m + \frac{k_t \cdot k_s}{k_t + k_s}\right)$$

$$F_{ma} = x_a \cdot \left[\left(\frac{k_s + k_t}{k_t}\right) \cdot \frac{k_m}{k_t} \cdot \frac{k_t \cdot k_s}{k_t + k_s} + \left(\frac{k_s + k_t}{k_t}\right) \cdot k_m + \left(\frac{k_s + k_t}{k_t}\right) \cdot \frac{k_t \cdot k_s}{k_t + k_s}\right] = x_a \cdot \left[\frac{k_m \cdot k_s}{k_t} + k_s \cdot \frac{k_m}{k_t} + k_t \cdot \frac{k_m}{k_t} + k_s\right] = x_a \cdot \left[2 \cdot \frac{k_m \cdot k_s}{k_t} + k_m + k_s\right]$$

### MUSCLE FORCE EQUATIONS

$$\mathbf{F_{ma} = x_a \cdot \left[2 \cdot \frac{k_m \cdot k_s}{k_t} + k_m + k_s\right]}$$

***INTERNAL MUSCLE FORCE***

$$\mathbf{F_{me} = k_s \cdot x_a}$$

***EXERTED MUSCLE FORCE TRANSMITTED TO SKELETON***

$$\mathbf{\frac{F_{me}}{F_{ma}} = \frac{k_s}{k_m + 2 \cdot \frac{k_m \cdot k_s}{k_t} + k_s}}$$

***RATIO OF TRANSMITTED MUSCLE FORCE TO INTERNAL MUSCLE FORCE***

**Figure S2. Derivation of muscle force equations.** Modeling the muscle, tendons, and skeleton as linear elastic springs enables deriving an expression for muscle internal actuation force and external transmitted force as a function of muscle, tendon, and skeleton stiffnesses.

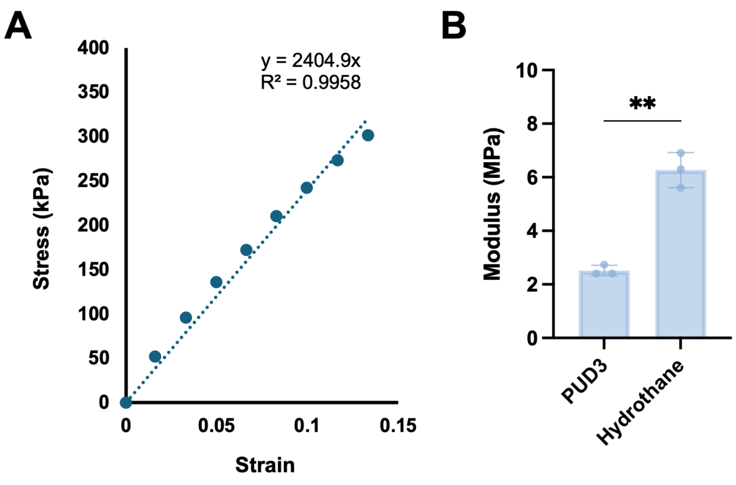


**Figure S3. Tendon mechanical testing.** (**A**) Representative data from tensile testing of a PVA:PAA hydrogel tendon with PUD3 backing. (**B**) Elastic modulus for tendons with PUD3 and Hydrothane backings (n = 3 per group, Welch's t-test, ** $p < 0.01$).

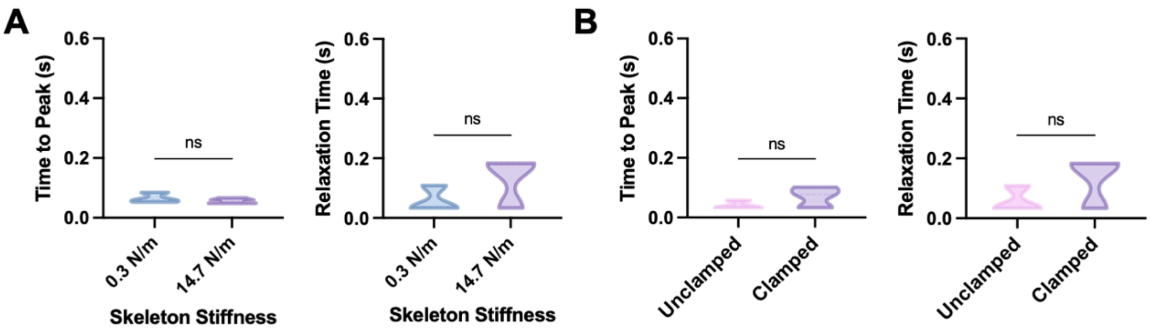


**Figure S4. Contractile dynamics of MTUs.** (**A**) Time-to-peak-force (left) and relaxation time (right) of MTUs on compliant flexure skeletons (0.3 N/m) and stiff gripper flexure skeletons (14.7 N/m). (**B**) Time-to-peak-force (left) and relaxation time (right) of MTUs on stiff gripper flexure skeletons (14.7 N/m) in unclamped and clamped configurations.

**Video S1. Impact of MTU pre-stretch on contractile displacement.**

**Video S2. Impact of stimulation frequency on MTU contractile displacement.**

**Video S3. MTU fatigue over 30 minutes at 4 Hz stimulation.**

**Video S4. MTUs on stiff gripper skeletons.**
